# Chromosomal Characteristics of Salt Stress Heritable Gene Expression in the Rice Genome

**DOI:** 10.1101/2021.01.05.425484

**Authors:** Matthew T. McGowan, Zhiwu Zhang, Stephen P. Ficklin

## Abstract

**Background:** Gene expression is potentially an important heritable quantitative trait that mediates between genetic variation and higher-level complex phenotypes through time and condition-dependent regulatory interactions. Increasing quantities of high-throughput DNA and RNA sequencing and standardization of research populations has resulted in the accumulation of overlapping -omics data allowing for deeper investigation into the genomic structure and conditional stability of gene expression traits. Therefore, we sought to explore both the genomic and condition-specific characteristics of gene expression heritability within the context of chromosomal structure, using a diverse, 84-line, *Oryza sativa* (rice) population under optimal and salt-stressed conditions.

**Results:** Heritability was estimated for the 84 genotypes with common tools and an approach using biological gene expression replicates similar to a twin study in humans. Overall, 5,936 genes were found to have heritable expression regardless of condition and 1,377 genes were found to have heritable expression only during salt stress. These genes with salt-specific heritable expression are enriched for functional terms associated with response to stimulus and transcription factor activity. Additionally, we discovered that highly and lowly expressed genes, and genes with heritable expression are distributed differently along the chromosomes in patterns that follow previously identified chromosomal conformation capture (Hi-C) A/B chromatin compartments. Furthermore, multiple genomic hot-spots enriched for genes with salt-specific heritability were identified on chromosomes 1, 4, 6, and 8. These hotspots were found to contain genes functionally enriched for transcriptional regulation and overlaps with a previously identified major QTL for salt-tolerance in rice.

**Conclusions:** Investigating the heritability of traits, and in-particular gene expression traits, is important towards a basic understanding of how regulatory networks behave across a population. Additionally, heritable gene expression architecture can be used for further exploration of gene-trait relationships, assist in interpretation and analysis of eQTLs, used as priors for approaches seeking to identification of potential biomarkers, or used in genomic selection modules with potential applications in plant breeding. This work provides insights into patterns of expression and spatial patterns at the chromosomal level.

## Background

Understanding the molecular mechanisms by which genetic variation influences complex quantitative traits remains a major goal of genetic research today. Current polygenic and omnigenic models posit that for complex traits, only a small proportion of heritable phenotypic variation can be explained by relatively few easily identified mutations with large effects. The remaining majority of heritable variation is due to a much larger quantity of low to moderate effect mutations. After more than a decade of research utilizing Genome-Wide Association Studies (GWAS) it is clear that many of these low to moderate effect genetic variants underlying complex traits tend to lie in regulatory regions of the genome rather than in protein coding regions. Furthermore, affected regions have been found to be enriched for genes that interact in highly interconnected regulatory networks [1]. Therefore, expression QTL (eQTL) studies seek to identify relationships between genetic variants and the genes on which they may have a regulatory effect by treating gene expression as the phenotypic trait for GWAS analysis.

The increasing number of studies investigating eQTLs in multiple plant species have revealed similar patterns of eQTL architectures. The location of eQTLs in relation to their affected gene are often referred to as *cis* and *trans* depending on whether they map respectively to the same relative location as the gene or elsewhere in the genome. While *cis* eQTLs tend to have larger effects on average compared to *trans* eQTLs, only a small proportion of genes appear to have *cis* eQTLs that explain a majority of their expression variance. Instead, many genes appear to have both *cis* and *trans* acting eQTLs with the most eQTLs being *trans* [2, 3]. Cross-gene eQTL analysis has revealed that many of these *trans* eQTLs are significantly enriched in genomic hotspots with wide reaching effects on gene expression [4, 5].

In any association study (GWAS or eQTL) characterization of heritability (*h^2^*) for the selected trait (e.g. phenotype or expression-level) is necessary to estimate genetic causality for the trait. Heritability is a fundamental genetics concept that describes how much of the variation in a given trait can be attributed to genetic variation [6]. It has demonstrated lasting usefulness in quantifying response to selection in plant breeding [7] and estimating disease risk in medicine [8]. Traditionally, heritability is estimated using known information about the genetic relationships between individuals. In human research, these known genetic relationships are usually in the form of monozygotic (identical) and dizygotic (fraternal) twins. In plant and animal research, pedigrees from controlled breeding populations are used to represent these genetic relationships. Another approach for estimating heritability uses high-density genotyping technologies such as single nucleotide polymorphism (SNP) arrays to infer genetic relationships. Genotype differences between individuals are used to calculate a genetic relationship matrix (GRM), also called a kinship matrix. This GRM is then used to estimate the proportion of phenotypic variance explained using linear mixed models. This approach is referred to as Genomic Relatedness Restricted Maximum Likelihood (GREML) and has multiple software implementations such as GCTA [9], EMMA [10], and rrBLUP [11]. Despite the large number of eQTL studies investigating gene expression, relatively few studies have explored genomic patterns of gene expression heritability using GREML-based estimates. Two studies in humans explored gene expression heritability of whole blood samples [12, 13], but similar research in plants is currently lacking.

Another area of gene expression research that is relatively unexplored is the influence of environmental factors. Even though differential gene expression (DGE) analysis is a highly active area of research, studies investigating variation in gene expression in response to environmental changes have primarily focused on condition, time, and tissue-specific expression variation. Yet these studies are limited to a few different genotypes, far below the necessary sample sizes required for performing eQTL analysis [14]. However, given that complex agronomic phenotypes are known to have significant genotype-by-environment interaction effects, exploring how these interactions affect gene expression variation may provide novel insights into the underlying architecture of these phenotypes.

An important consideration prior to exploration of heritability is understanding any potential bias from variation that underlies the bimodal distribution of gene expression. It has been shown that gene expression when quantified with RNA-seq data has a bimodal structure such that lowly expressed (LE) genes and highly expressed (HE) genes appear as two overlapping distributions with LE genes centered in the negative log2 range and the other in the positive log2 range [15]. The source of this bimodality is a currently a topic of debate. One theory suggests the lower distribution is due to an unknown combination of transcriptional noise, ambiguous read mapping, contamination, cell type heterogeneity, and sequencing errors. Thus, many only use the HE genes for downstream research [16]. However, there is evidence that transcripts from the low abundance distribution are transcribed mRNA and not artifacts or small RNA molecules [17].

Another consideration for exploration of gene expression heritability, related to non-normal gene expression distributions, is that transcriptional repression has been shown to be correlated with the 3D conformational structure of chromosomes in the nucleus including chromatin and centromeric structures [18]. Chromatin alteration in plants has been shown to play important roles in tissue-specific specialization [19, 20], stress response [21–23], and suppression of transposable elements [24, 25]. Plant genomes have been found to possess active and repressive genome territories referred to as the A and B compartments which correspond to euchromatic and heterochromatic regions, respectively [26, 27]. While these compartments have been found to be largely stable across tissues, it remains unclear how stable these compartments are across changing environmental conditions known to alter chromatin states such as abiotic stress.

In this study, we sought to address the limitations and considerations just described for gene expression heritability by exploring the 2D and 3D chromosomal characteristics of heritable gene expression using an RNA-seq dataset of 84 individuals of the *Oryza sativa* Rice Diversity Panel 1 (RDP1) previously reported [28]. We explored patterns of missing values in the RNA-seq data (i.e. missingness) and the distribution of highly expressed (HE) and lowly expressed (LE) genes across the 2D chromosomal structure. Heritability was calculated independently for salt stress and control conditions and their distribution was also explored across the 2D genomic structure. We then explored the relationship of HE and LE genes to the Hi-C analysis of rice chromatin structures.

## Results

### Gene Expression

For the 55,986 annotated gene transcripts in the Michigan State University (MSU) v7.0 Oryza sativa Nipponbare (rice) assembly [29], the distribution of missing values (genes with no measured expression) followed a U-shaped distribution with most genes having either a high or low missing rate and relatively few genes having moderate levels of missingness. We classified genes as having constitutive, mixed, or repressed expression patterns if non-zero expression was observed in >95%, 5-95%, or <5% of samples, respectively (Figure 1a). Overall, non-zero gene expression followed a clear bi-modal distribution consisting of a mode of highly expressed (HE) genes with positive log_2_ TPMs and a second mode of lowly-expressed (LE) genes with negative log_2_ TPMs (Figure 1b). Genes with constitutive expression occupied the HE mode, while genes with a mixed or repressed expression pattern matched the LE mode. Thus, HE genes are both highly expressed and highly present (few missing values) while LE genes are lowly expressed and lowly present. Furthermore, genes rarely changed from constitutive, mixed or repressed between salt or control treatments (Table 1). While there were a small number of genes that switched categories between conditions, there were no genes that changed from constitutive to repressed.

**Figure 1.**
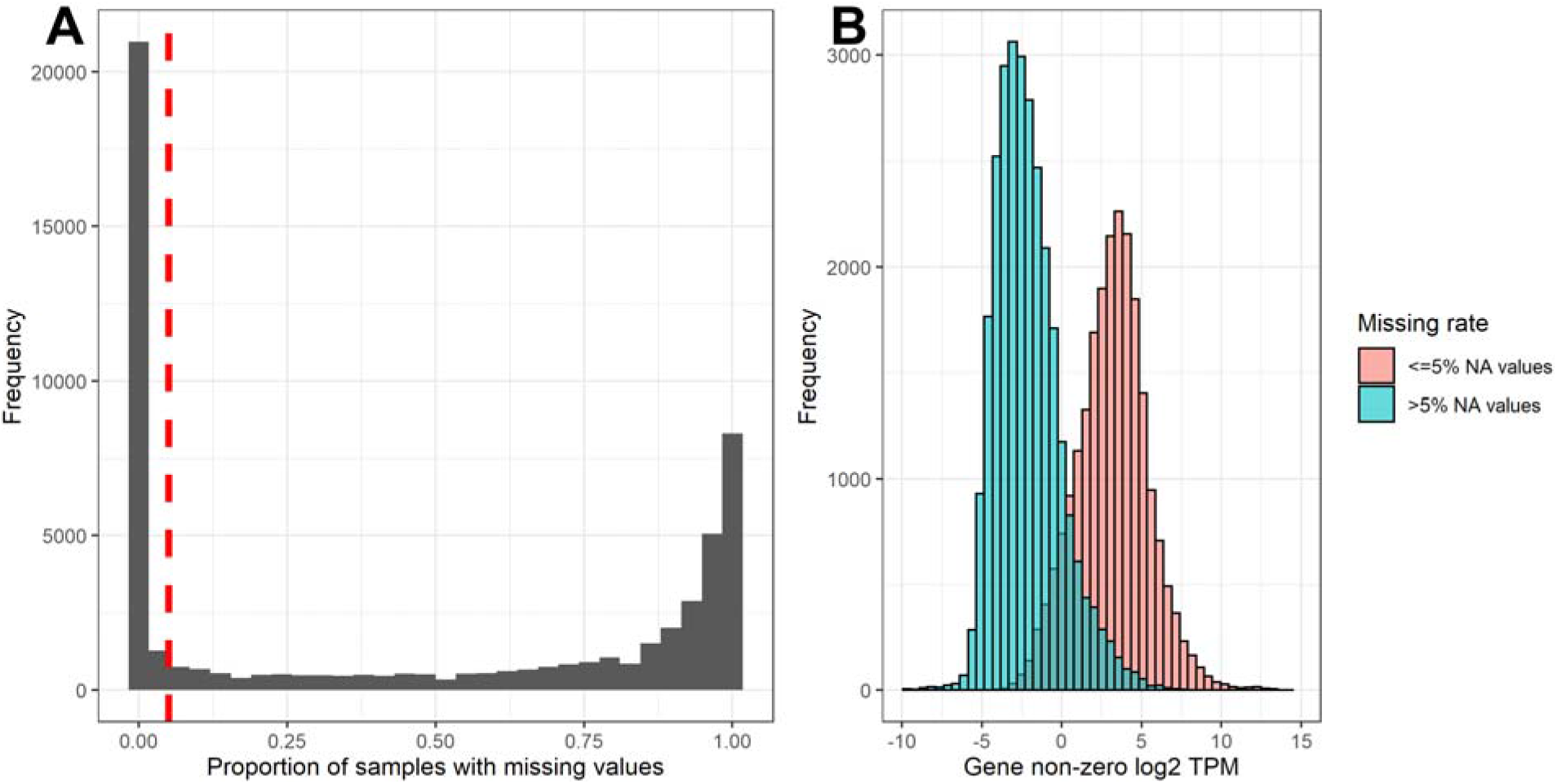
Bimodal Gene Expression Patterns: The proportion of samples with missing values was calculated for each gene. The overall distribution of this missing rate (A) is bimodal with the majority of genes either having few (<5%) or many (>95%) missing values. Genes were classified as ‘constitutive’ (<5% missing), mixed (5-95% missing), or repressed (>95% missing). The mean value of non-zero TPMs for expressed genes also had a bimodal distribution based on the missing rate (B).

**Table 1.** Cross-conditional Expression-Level Categories

|  |  | Salt-stress |  |  | Totals |
| --- | --- | --- | --- | --- | --- |
|  |  | Constitutive | Mixed | Repressed |  |
| Control | Constitutive | 16372 | 363 | 0 | 16735 |
|  | Mixed | 91 | 25116 | 932 | 26139 |
|  | Repressed | 0 | 1007 | 12105 | 13112 |
|  | Totals | 16463 | 26486 | 13037 |  |

### Heritability

#### Comparison of heritability results

Correlation of gene expression biological replicates on a per-gene basis was calculated as a potential estimate for heritability, similar to twin-based measures of heritability in humans. Replicate heritability values were then compared to both GREML estimates of heritability using a genotypic mean (two-step) and GREML estimates that included replication as a random effect in the model.

Due to the relatively small sample size, there were many genes where the GREML heritability (single-step or two-step) could not be reliably predicted with a mixed linear model resulting in an inflated number of genes with low heritability estimates (0-0.2) and a wide 95% confidence interval (Additional File 1, Fig. S1). There was strong correlation between replicate heritability versus single-step GREML (⍰ = 0.89), indicating that gene expression heritability can be estimated using the biological replicates expression data. However, the correlation of the two-step method was moderate when compared to the one-step approach (⍰ = 0.41) and with replicate heritability approach (⍰ = 0.45) (Figure 2). Results in Figure 2 are for the control condition, but patterns were similar for the salt condition (Additional File 1, Fig. S2).

**Figure 2.**
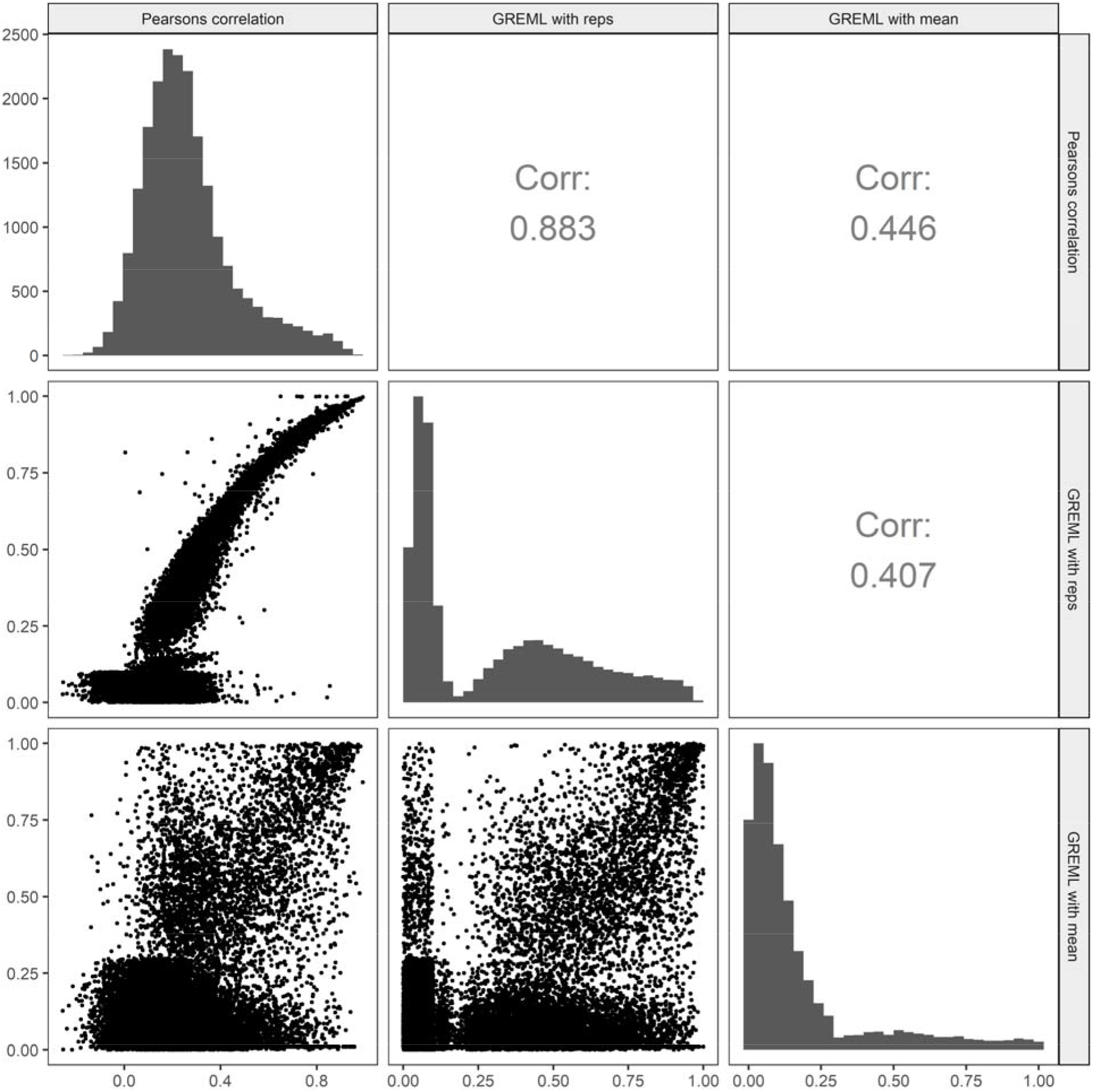
Comparison of Heritability Calculation Methods for the Control condition: Pairwise correlation between repeatability (Pearsons), single-step GREML (with replicates), and two-step GREML (using the genotypic mean) for the control condition

#### Condition-specific heritability classification

To identify a significance threshold for expression heritability, randomized permutation tests of shuffled gene expression values were used to calculate a null heritability distribution. Using this null-distribution, a significance threshold was calculated using a fixed type-I error rate (◻<= 0.01) (Figure 3a). Genes were classified whether they were significantly heritable for control and salt-stress conditions (Figure 3b). While most genes with heritable expression appeared to have conserved heritability for both control and salt-stress conditions (*n*=6,851), there were a considerable number of genes significantly heritable only during control (n=3,599) or salt-stress (*n*=1,377). These genes with condition-specific heritability were less heritable than genes that were heritable across both conditions (Additional File 1, Fig. S3). Genes heritable in both salt stress and control were correlated symmetrically along the diagonal (Figure 3b), indicating no condition-specific bias.

**Figure 3.**
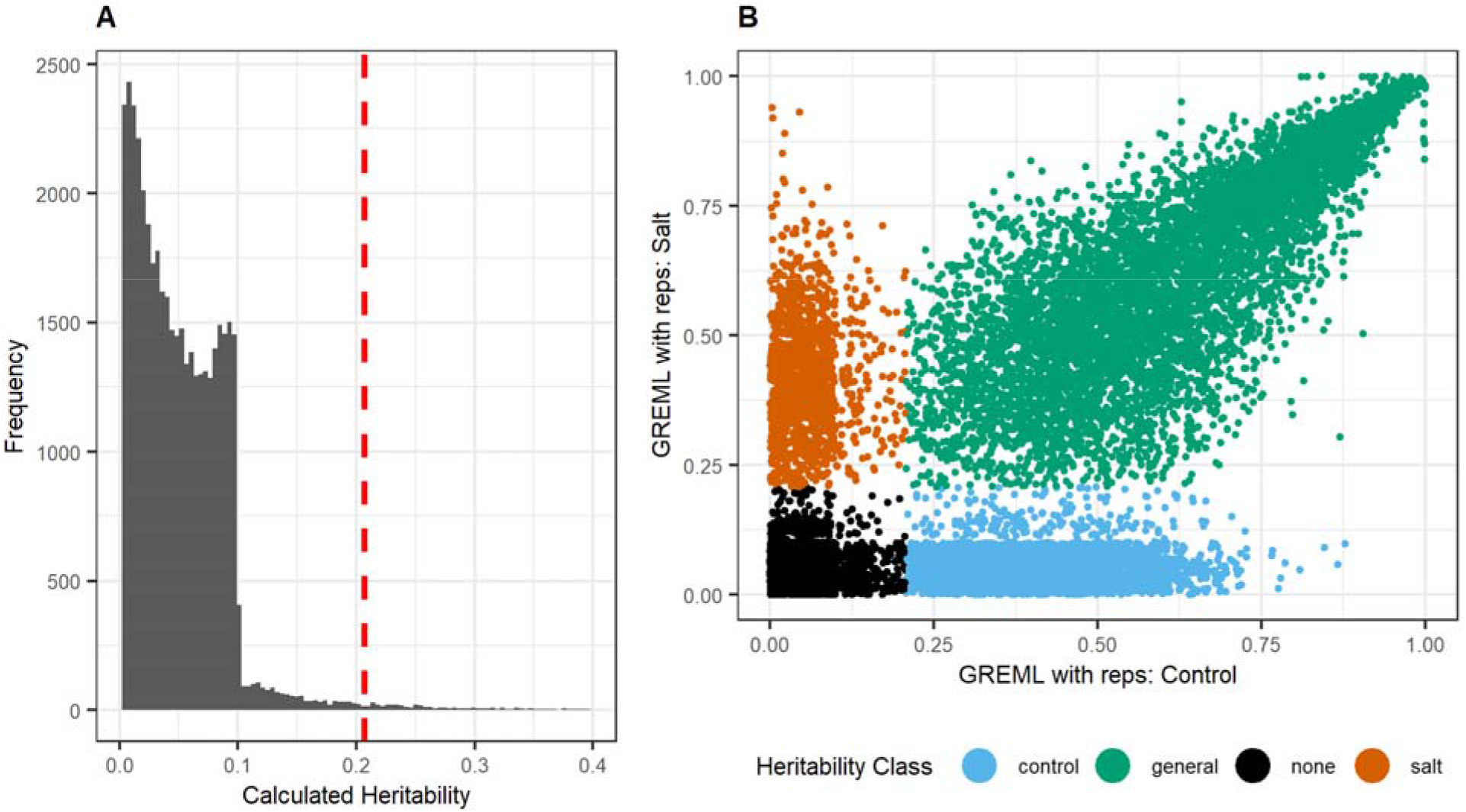
Classification of gene expression heritability. Randomly shuffled gene expression values were used to determine a null-distribution of heritability estimates for genes with non-heritable expression (A). The dashed red line indicates the quantile for a fixed type-1 error (◻=0.01). This quantile threshold was used to classify each gene as having significant heritability in salt treatment, control or both (B).

### Chromosomal Structure and Conformation

#### HE and LE genes follow distinct 2D spatial patterns

The spatial distribution of constitutive, mixed, and repressed genes was visualized along the chromosomes using a sliding window of 3Mb at 100Kb intervals. Empirically, constitutive genes appear enriched on the ends of chromosomes and depleted near pericentromeric regions (Figure 4). For metacentric chromosomes, this pattern formed a U-shape centered on the centromere. Densities for genes with repressed and mixed expression were often inverse of constitutive genes and appear enriched near the centromere and depleted at the chromosome ends. Reductions in density of constitutive genes were not always centered on the centromeric regions. For example, subtelocentric chromosomes 4, 9, and 10 (and chromosome 11 to a lesser extent) show this asymmetry as the short chromosomal arms appeared relatively devoid of genes with constitutive expression (Figure 4).

**Figure 4.**
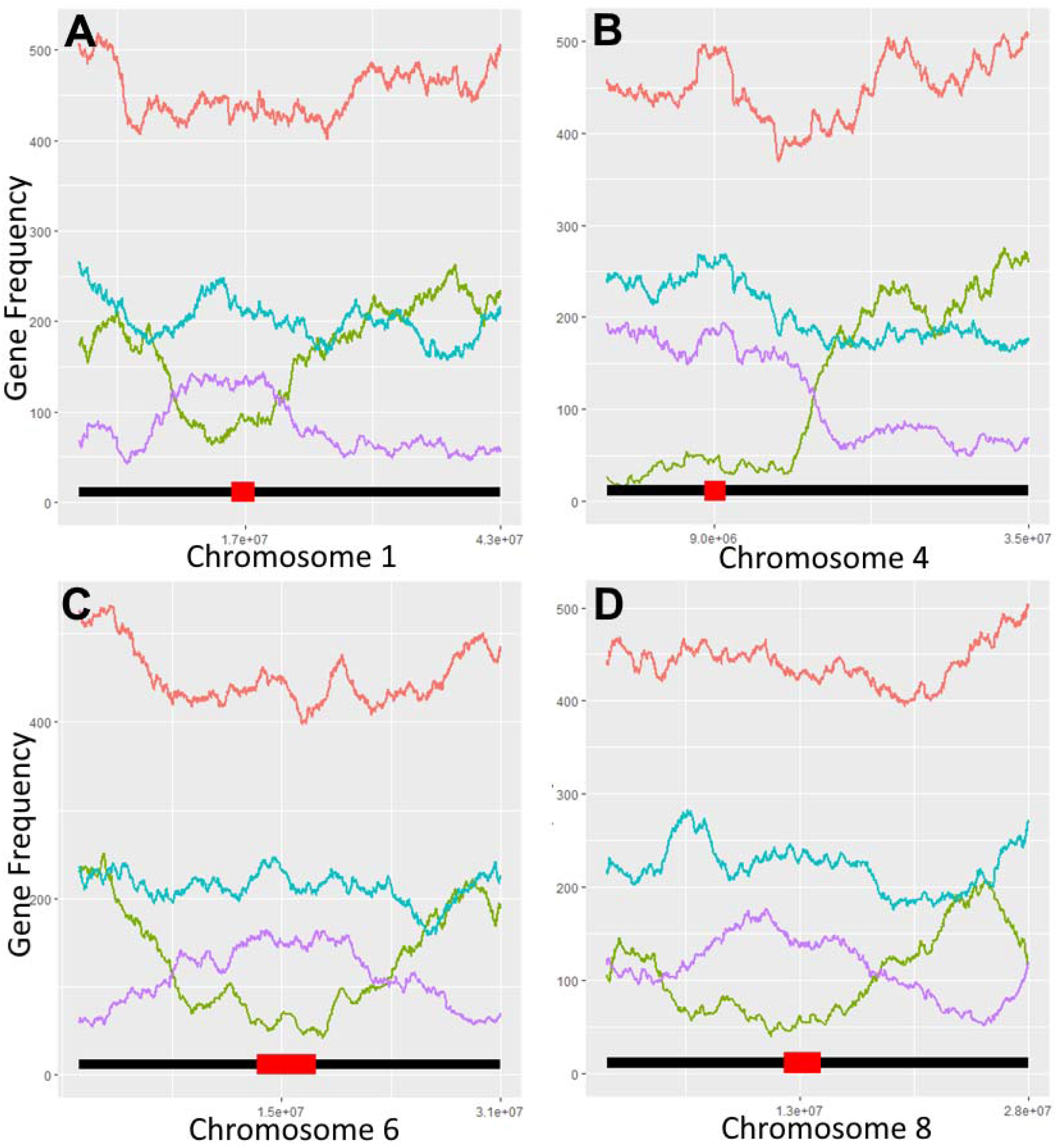
Gene density distributions across chromosomes 1 (A), 4 (B), 6 (C), and 8 (D). Pericentromeric regions are indicated by red boxes at the bottom of each density plot. Overall gene annotations appear roughly uniform across each chromosome (red). Genes with constitutive expression (>95% samples) of samples are enriched on the distal ends of chromosome arms and depleted near pericentromeric regions (green). Genes with repressed expression (<5% of samples) are enriched near pericentromeric regions (blue). Genes with mixed expression (5-95% of samples) largely follow the same distribution as repressed genes (pink).

#### Comparison of gene expression and HI-C A/B chromatin compartments

Regarding 3D characteristics of expressed genes, densities of genes (when calculated using a fixed 100kb window size) were highly correlated (⍰ = 0.7-0.9) with A/B chromatin compartments identified with the first principal component of PCA analysis of a Hi-C contact map [27] (Additional File 1, Figs. S4-S6). Euchromatic A compartments corresponded to genes that were constitutively expressed across all genotypes. Conversely, heterochromatic B corresponded to genes with either mixed or repressed expression across genotypes.

### Salt-specific spatial enrichment analysis

When the spatial distribution of genes with salt-specific heritability was compared to the distribution of genes with non-specific heritability, 22 windows were identified on chromosomes 1, 4, 6, and 8 that passed a permutation-based *p*-value threshold (◻=0.01) (Figure 5, Table 2). This test indicates where the genome is enriched for salt-stress specific expression. Other chromosomes did not have significantly enriched windows (Additional File 1, Figs. S7-S9). Adjacent and overlapping windows were combined into five contiguous regions. Gene ontology enrichment analysis of heritable genes in these regions identified terms of transcription factor activity (GO:0003700), response to endogenous stimulus (GO:0009719), nucleic acid binding (GO:0003676), and DNA binding (GO:0003677) (Additional File 2). When compared to previous GWAS studies, there were overlaps between these regions and QTLs identified for salt-tolerance related traits. In particular, a 3 Mb window on chromosome 4 directly overlaps with a highly significant 575 Kb QTL identified from a previous GWAS that used the same RDP1 panel that was significant for sodium and potassium accumulation in root tissue [28]. Fine mapping of this QTL identified HKT1;1, a sodium-transporter gene (LOC_Os04g51820) that is the likely causal gene. It was also determined that altering the expression of this gene using RNA-interference lines significantly affected both shoot and root growth under saline conditions [28].

**Figure 5.**
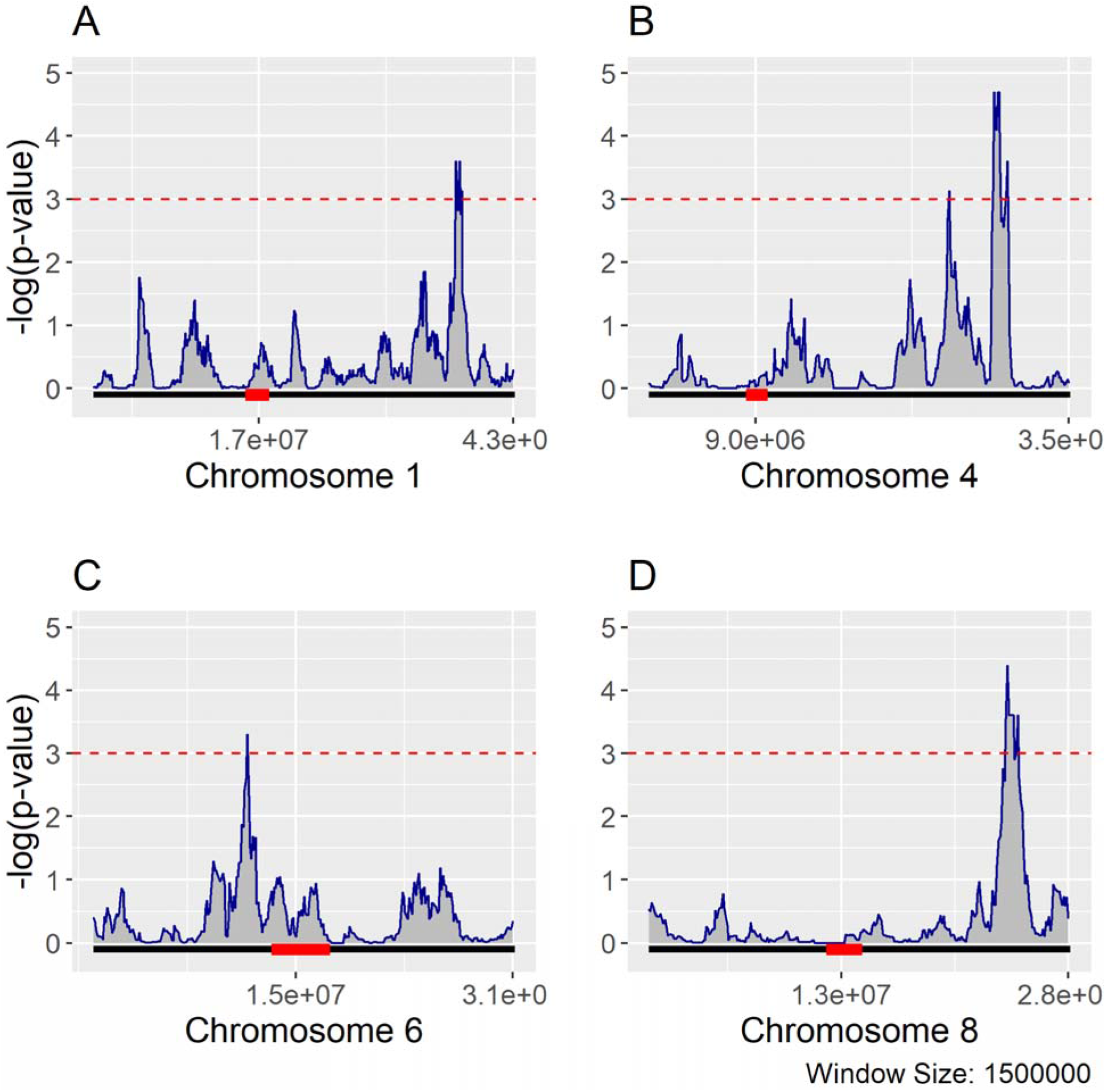
Salt-specific Heritable Gene Enrichment. Genome enrichment for salt-specific heritable expression: Using a sliding window size of 1.5 Mb at 100 Kb intervals, chromosomes were tested for enrichment of genes with salt-specific heritability using all genes with heritable expression (salt-specific, optimal-specific, and general) as the null distribution. P-values were adjusted for multiple-testing using a permutation based approach. Using a critical value of 0.001 (red dotted line), significant windows enriched for salt-specific heritability were identified on chromosomes 1, 4, 6, and 8. Centromeres are located with a red box.

**Table 2.** Genome windows enriched for salt-specific heritable expression.

| Chromosome | Start<br>Position | End<br>Position | Heritable<br>genes | Fisher's test<br>adjusted *p*-value |
| --- | --- | --- | --- | --- |
| 1 | 36,450,000 | 38,550,000 | 19 | 2.5E-04 |
| 4 | 24,550,000 | 26,050,000 | 17 | 7.5E-04 |
| 4 | 28,250,000 | 30,950,000 | 19 | 2.0E-05 |
| 6 | 10,650,000 | 12,150,000 | 13 | 5.0E-04 |
| 8 | 23,350,000 | 25,650,000 | 17 | 4.0E-05 |

In summary, results show missingness is the cause of bimodality in the salt-stress gene expression data. Regarding 2D characteristics, HE and LE genes have distinct distribution patterns in relation to the centromeric location of the chromosomes. Additionally, salt-specific heritable genes follow similar 2D distribution patterns but are also highly correlated with 3D conformation following Hi-C identified A/B compartments. We also identified several significant genomic hot-spots enriched for genes with salt-specific heritability on chromosomes 4 which is concordant with previous GWAS studies investigating salt tolerance phenotypes in a similar population as well as 3 additional windows on chromosomes 1, 6, and 8.

## Discussion

### Gene Expression

It has been suggested that low abundance mRNA identified in the LE distribution of TPM values may not be transcribed into proteins. Comparisons between lowly abundant genes in human metazoan cells and proteome quantification in human embryonic cells did not indicate that LE genes are translated [17]. While the results presented here do not definitively answer the question, the patterns observed both in the bimodal distribution (Figure 1) and the cross-conditional table (Table 1) provide insight regarding variation of transcriptional repression. Genes with few missing values tend to have high TPM expression values. However, when a gene had a zero value, in any sample, then most non-zero values were in the LE distribution. Furthermore, when these patterns were compared between salt and control conditions, there were no genes that switched from repressed expression to constitutive expression in the population. Considering that four times as many genes shifted between mixed and repressed states (1,939 genes) compared to genes that shifted between mixed and constitutive states (454 genes), one explanation is that many of these genes are located within chromosomal regions that are still largely repressed, but that this repression is incomplete and a low level of transcription still occurs. However, it is also possible that some of these conditional lowly expressed genes are being translated into proteins. Given that RNA-seq samples in this experiment consisted of homogenized shoot samples containing multiple cell types, cell-type specific expression could also explain genes that are lowly expressed. While the sample size was too small to reliably calculate the heritability of mixed and repressed gene expression using logistic models, PCA of the gene expression matrix encoded as ordinal zero, low, or high expression suggests that there is a large amount of additional transcriptional variance that closely matches the genotypic population structure (Additional File 1, Fig. S10). This variation may not be captured in current RNA-seq approaches that only consider TPMs from the HE distribution such as differential expression models using the negative-binomial distribution.

Regarding the notion that LE genes are not translated into proteins, this assumption is based on limited evidence that compared different cell types in different conditions. However, it may be too early to rule out potential translation of LE genes. Plant genomes have been shown to undergo drastic heterochromatic reorganization in response to abiotic stimuli including salt-stress [30]. The high correlation between LE genes and heterochromatic regions of the genome may suggest that rather than being untranslated, the low expression of these genes could be related to cell type or condition-specific responses, which would lead to their proteins not being observed in previous proteomics studies that used different conditions and genotypes.

### Heritability

The importance of using biological replicates for differential gene expression analysis has already been explored [31, 32] but this research also indicates that biological replicates provide important information for models estimating gene expression heritability. Considering the inherent noise that can be introduced by natural variation in gene expression such as circadian rhythm, the inclusion of biological replicates should be considered an indispensable aspect of RNA-seq experimental design. Previous research investigating the statistical power of RNA-seq based differential expression analysis indicated that at least six biological replicates were required to identify the majority of differentially expressed [32]. However, no studies have explored how increasing the number of biological replicates can improve the power of models that estimate gene expression heritability. Considering that these models can also benefit from increasing the number of genotypes, there is need for quantifying the power trade-off between the number of genotypes and the number of biological replicates for accurately estimating gene expression heritability.

Another result of interest is that the two-step GREML showed only moderate correlation with both the replicate-based and one-step GREML estimates. Differences in how genetic effects are distributed may explain this. Previous reports on eQTLs underlying gene expression heritability in humans suggest that highly heritable gene expression tends to be controlled by relatively few *cis* eQTLs with strong, non-additive, effects [33, 34]. Conversely, heritable complex traits and moderately heritable gene expression tend to be controlled by many small additive effect mutations [35, 36]. This difference in how genetic effects are distributed may explain why GREML heritability estimates using mean expression was only lowly correlated with repeatability. Previous studies investigating heritability in human populations (with a much larger sample size than this study) split markers into separate *cis* and *trans* components in the GREML model where the *cis* random effects only included markers surrounding the gene being tested with the remaining markers included in the model as a separate *trans* random effect [13]. The approach for splitting *cis* and *trans* components in these studies used only markers within a 1 Mb fixed window around a gene as the *cis* component (that was likely to capture any promoter regions) and treated all other markers as a separate *trans* component. The purpose for this is that mutations near the coding sequence and surrounding promoters seem more likely to have large effects on gene expression and thus would follow a different underlying distribution of effect sizes compared to mutations occurring elsewhere in the genome. In these human studies, the average overall mean heritabilities were reported to be between 0.15 and 0.26 with the proportion of heritability explained by *cis* markers ranging from 20-40% depending on the tissue and population studied. A smaller microarray-based eQTL study in an *A. thaliana* RIL population reported a similar heritability distribution [2]. Notably, they also observed many genes that exhibited transgressive segregation and suggested that nonadditive genetic variation may be significantly contributing to overall expression heritability in plants.

The sample size of the data used in this study was too low to reasonably split markers into separate *cis* and *trans* random effects in the additive GREML model to allow for direct comparison to previous studies. However, the low correlation between the two-step GREML additive-only model and the one-step GREML model that included replicates as a random effect supports the idea that gene expression traits have a genomic architecture that cannot be captured well by treating all genome-wide markers as a single additive random effect distribution. One possible alternative for modeling gene expression traits that could avoid an arbitrary fixed window for splitting markers into *cis* and *trans* components is to use variable selection methods that can accommodate mixed distributions of marker effects. There is considerable similarity between the previously used strategy of modeling separate *cis* and *trans* components and Bayesian models used for genomic selection which can accommodate many different prior distribution assumptions [37]. However, challenges remain for testing whether these Bayesian methods can more effectively estimate marker effects underlying transcriptome-wide gene expression. First, there are many different prior distributions proposed for performing Bayesian genomic selection and selecting a suitable prior distribution is non-trivial considering that the underlying architectures of heritable gene expression are heterogeneous [38, 39]. Secondly, even with parallelization, the Markov chain Monte Carlo (MCMC) algorithms involved have considerably higher computational costs compared to GREML making intensive testing difficult.

### Chromosomal Structure and Conformation

The strong correlation between gene expression chromosome densities and HiC compartment predictions supports the paradigm that pericentromeric regions play an important transcriptional regulatory role in the 3D conformation of chromosomes in the nucleus and primarily correspond to heterochromatic B compartments in rice. For example, HE genes with constitutive expression patterns are more likely to be located in euchromatic A compartments, while LE genes with low and repressed expression are more likely to be located in heterochromatic B compartments. Therefore, the strong relationship identified between a gene’s expression pattern and its position in the chromosome may have important implications for predicting the effects of structural variations such as translocation or gene duplication events. Such an understanding may improve studies exploring the role of duplicated genes, as it may be essential to consider where in the chromosome duplicate genes are located and how the surrounding regulatory landscape is different (such as a shift in chromatin compartment).

### Overlap Between Salt Stress QTLs and Expression Heritability

An interesting observation regarding the overlap between salt-tolerance associated QTLs identified in the RDP1 population using GWAS and the windows enriched for salt-stress specific heritable expression is that the current putative causal gene underlying the largest salt-tolerance QTL in this population, OsHKT1;1 (LOC_Os04g51820), did not exhibit heritable gene expression after accounting for population structure. However, many genes within close proximity to this gene did have heritable expression and this region was particularly enriched for salt-specific expression heritability. This indicates that causal genes underlying complex phenotypes may have indirect effects on gene networks. One possible explanation for this is that genes that co-participate in shared biological pathways have been shown to cluster in the same chromosomal region [40]. However, this clustering does not occur in all plant pathways and there are currently many theories for why some pathways are genomically clustered and others are not [41]. One of these theories is the ‘coinheritance argument’ where genetic linkage of genes with shared roles in a complex trait can promote the accumulation of favorable genes and reduce risk of disruption via recombination. Given that salt-tolerance is a trait in rice with a history of both evolutionary and artificial selection, this theory may explain the clustering observed.

### Implications

Results show that the relatively small sample sizes in this study (compared to typical GWAS studies) were able to identify regions of the genome enriched for condition-specific heritable gene expression. This approach could be used to identify genes involved with conditional transcriptomic plasticity. Identifying heritable genes with genotype-by-environment specific behaviors may be useful to breeders in MAS approaches to select for mutations with more isolated trait-specific effects, across genotypes, and avoid the selection of mutations with strong epistatic effects.

While it is generally accepted that the genome-wide distribution of marker effects for complex traits is non-uniform, there are few approaches for determining how non-uniformity relates to the physical genome. However, the chromosome-level patterns of gene expression heritability observed in this study could potentially be used as prior estimates of possible marker effect distributions for Bayesian genomic selection models. Even if the underlying true distribution may have cryptic condition-specific components outside the scope of available RNA-seq data, a large proportion of heritable expression was observed for both conditions. For example, there were multiple regions of the genome with relatively few genes with heritable gene expression for either condition. Markers within these regions could be assigned low prior probabilities of having strong effects. In contrast, we also identified regions of the genome with high general and condition-specific heritable expression. Markers within this region could be assigned higher prior weights, especially when they are located in trait related conditional hotspots.

## Future Considerations

As the increasing number of studies in plants utilizing standardized genetic diversity panels for producing -omics based data allow for rich multi-dimensional research into biological systems, results observed in this study provide suggestions for future research. While further experiments investigating these hotspots for salt-specific heritable expression are required for validation, results regarding missing values and their relation to bimodal expression patterns highlight the need for more overlapping -omics data. First, use of larger genotype panels for transcriptomic sequencing with more biological replications would improve heritability estimates and support greater exploration of *trans* effects. Second, access to high-resolution contact maps would allow for further investigation into the roles lower-level chromatin structures (such as topologically association domains) play in regulatory variation for how plants respond to stress. While many RNA-seq experiments primarily focus on analyzing highly expressed genes, this research indicates that genes with low non-zero expression also have distinct spatial patterns that may provide evolutionary value and should be further explored. The addition of conditionally matched proteomics data would help resolve the open question if any of these lowly-expressed genes are ever translated into proteins.

## Methods

### Genotype Data

All rice accessions used in this research are from the Rice Diversity Panel 1. This panel consists of 421 purified, homozygous rice accessions that include both landraces and elite rice cultivars worldwide. Genotypes for the entire panel were obtained from the online project repository for the Rice Diversity Project [42]. In particular, this research used a set of 44k SNPs obtained from a combination of array and sequencing-based approaches. Missing genotypes were imputed using LD-kNNi [43]. The cross-validated accuracy using known genotypes was found to be highly accurate (R^2^ = 0.98). Markers with an imputed minor allele frequency of less than 5% were removed leaving a total of 31,374 markers for further analysis.

### Gene Expression Data

RNA-seq sequence files for a subset of rice accessions (*n*=92) from the RDP1 panel were identified and sourced from the National Center for Biotechnology Information (NCBI) sequence read archive (SRA) listed under Gene Expression Omnibus (GEO) project GSE98455. This previously published data originates from a project investigating salt-stress related gene co-expression network modules [28]. Briefly, seedlings of each accession were subjected to either optimal or salt-stress conditions for 24 hours and afterwards, shoot-tissue RNA was extracted and sequenced. Each treatment has two biological replicates originating from separate but genetically identical inbred accessions for a total of 368 RNA-seq samples. Only accessions that had replicates for both conditions were used (*n* = 84) for a filtered total of 336 samples.

RNA-seq files were downloaded and processed using the GEMmaker v1.1 pipeline for gene expression analysis [44]. This pipeline streamlines the process of calculating a gene expression matrix (GEM) from large numbers of raw FASTQ [45] sequencing files. GEMmaker was configured to download the GEO project GSE98455 sequence files using the SRA toolkit [46], perform quality control with FastQC [47] and quantify Transcripts-per-million (TPM) [48] expression values using Kallisto [49], a pseudo-alignment based tool. Gene annotations from the Michigan State University (MSU) Rice Genome Annotation Project (MSU release 7) were used for pseudo-alignment, which are based on the International Rice Genome Sequencing Project reference genome (Os-Nipponbare-Reference-IRGSP-1.0) [50]. TPM values were calculated at the gene level rather than the isoform level due to limited annotation of alternative splicing in rice. TPM values were log2 transformed. The sample and gene-wise distributions of mean log2 TPM and proportion of missing values were assessed.

### Structural analysis

Prior research on this population’s structure indicated that the panel has five major sub-groups [42]. We replicated the structural analysis with the subset of RDP1 individuals used in this study and found the same conclusion. Based on principal-component analysis (PCA), the top three components were found to capture a majority of genetic variance across subgroups (61%) (Additional File 1, Fig. S11). Initial inspection of pairwise TPMs indicated the presence of population structure matching the major classes of rice identified from marker-based principal component analysis. Because of the high collinearity between markers separating these groups and clusters of gene expression, not accounting for this population structure could lead to inflated heritability estimates [51]. For linear mixed models, this can be addressed by including subpopulation identifiers or principal components as fixed effects within the model. However, for simpler repeatability metrics, such as Spearman correlation, this is not possible. Therefore, expression values were adjusted to remove the major subpopulation effects prior to calculating any heritability estimates. This was done by fitting the expression values for each gene with a linear model including the top three principal components calculated from the genotype matrix as independent fixed effects. The remaining residuals were then used as adjusted gene expression values for calculating heritabilities for each gene. This adjustment was done separately for each condition and replicate. The distribution of the impact of structural adjustment indicates that while the repeatability of most genes slightly decreased when structure was removed, this decrease was larger for genes with clear clustering related to the population structure (Additional File 1, Fig. S12).

### Heritability Calculations

All statistics were performed using R 3.6.0 (R Core Team, 2018). Condition-specific heritabilities were calculated for the expression of each gene and condition using multiple methods. Genes with <5% missing values across all samples were used in heritability analysis. Heritability was first estimated using the similarity of expression between biological replicates. Because all accessions were inbred lines, and conditions were tightly controlled between lines, the Pearson or Spearman correlations between replicates within a shared condition were calculated as an upper-bound of the heritability of gene expression for that condition. Heritability was also calculated using single-step and two-step GREML algorithms implemented in the R ‘heritability’ package version 1.3 [52]. These GREML methods use a GRM calculated from genotypes to solve a linear mixed model for a quantitative phenotype with the efficient mixed-model association (EMMA) algorithm. This mixed model approach is commonly used for estimating heritability and genomic selection of agronomic phenotypes in crops [53]. The GRM was estimated using a standard method (VanRaden) implemented in the genetic ridge-regression R package ‘rrBLUP’ using version 4.6.1 [54]. Because expression values were already adjusted for subpopulation structure, principal components were not included in the GREML models. Default values were used for the convergence criterion (eps) and maximum iterations (max.iter). Two-step GREML calculation was performed by first calculating the genotypic expresssion mean across replicates and then regressing these mean values with a linear model treating kinship as a random effect using the *marker_h2_means* function. Single-step GREML was estimated using the *marker_h2* function but instead of regressing the genotypic mean, it includes replicate variance in the model as an added random effect [52]. The significance of heritability estimates compared to randomized gene expression were measured based on a one-tailed shuffled permutation test where the expression values for each gene were randomly shuffled for 40,000 iterations. This number of iterations was chosen based on hardware capabilities within a 48-hour window. Assuming that randomly shuffled gene expression vectors should not be heritable, the resulting heritability distribution of randomized expression was used to calculate a significance threshold based on a fixed type-1 error rate (◻=0.01). This threshold was then used to test whether each gene was significantly heritable (Figure 3a). Genes were then classified if they were significantly heritable under control, salt-stress, or both conditions (Figure 3b).

### Spatial enrichment analysis

The spatial distribution of expressed genes and their calculated heritabilities across each chromosome were compared between the control and stress conditions. This was done using a sliding window (1.5 Mb) and sampling interval to calculate the frequency or density of genes across each chromosome. Enrichment of salt-specific heritable genes for each window was determined using a one-tailed Fisher’s exact test. Test *p*-values were adjusted for multiple testing bias by calculating a null-distribution for each window. First, random gene subsets of equivalent size to the number of salt-specific heritable genes were randomly drawn from all heritable genes. For each random subset, a Fisher’s test was performed for each window resulting in a window-specific *p*-value. This process was bootstrapped for 4,000 iterations (*n* = 4,000 and the resulting *p-*value distributions were used to calculate adjusted *p*-value quantiles. For windows where 4,000 iterations were not enough to assign a *p*-value quantile, these windows were further tested for up to 50,000 iterations until a stable quantile estimate was obtained. Genes significantly enriched for salt-specific heritable expression within windows were tested for functional term enrichment using the Comprehensive Annotation of Rice Multi-Omics (CARMO) tool [55].

### Multi-omics Integration

The spatial distribution of gene expression heritability was then overlaid with other types of - omics data. The densities of genes with different expression patterns (constitutive, mixed and repressed) were tested for correlation with chromatin A/B compartment eigenvectors from previously published Hi-C analysis [27]. The Hi-C analysis used a fixed bin width of 100kb. This fixed window size was then used to calculate the gene densities of different expression patterns and correlated to the A/B eigenvectors.

## Data availability

A repository containing R scripts for replicating heritability analysis and visualization of results are available on the Open Science Framework [56].

## Acknowledgments

We thank Lei Gong and staff at the Key Laboratory of Molecular Epigenetics of the Ministry of Education at Northeast Normal University, Changchun, China for providing their Hi-C A/B compartment eigenvalue data.

## Contributions

Analyzed the data: MM. Wrote the paper: ZZ MM SF. All authors have read and approved the manuscript.

## Ethics declarations

### Ethics approval and consent to participate

Not applicable

### Consent for publication

Not applicable

