## Additional File 1 for "Chromosomal Characteristics of Salt Stress Heritable Gene Expression in the Rice Genome"

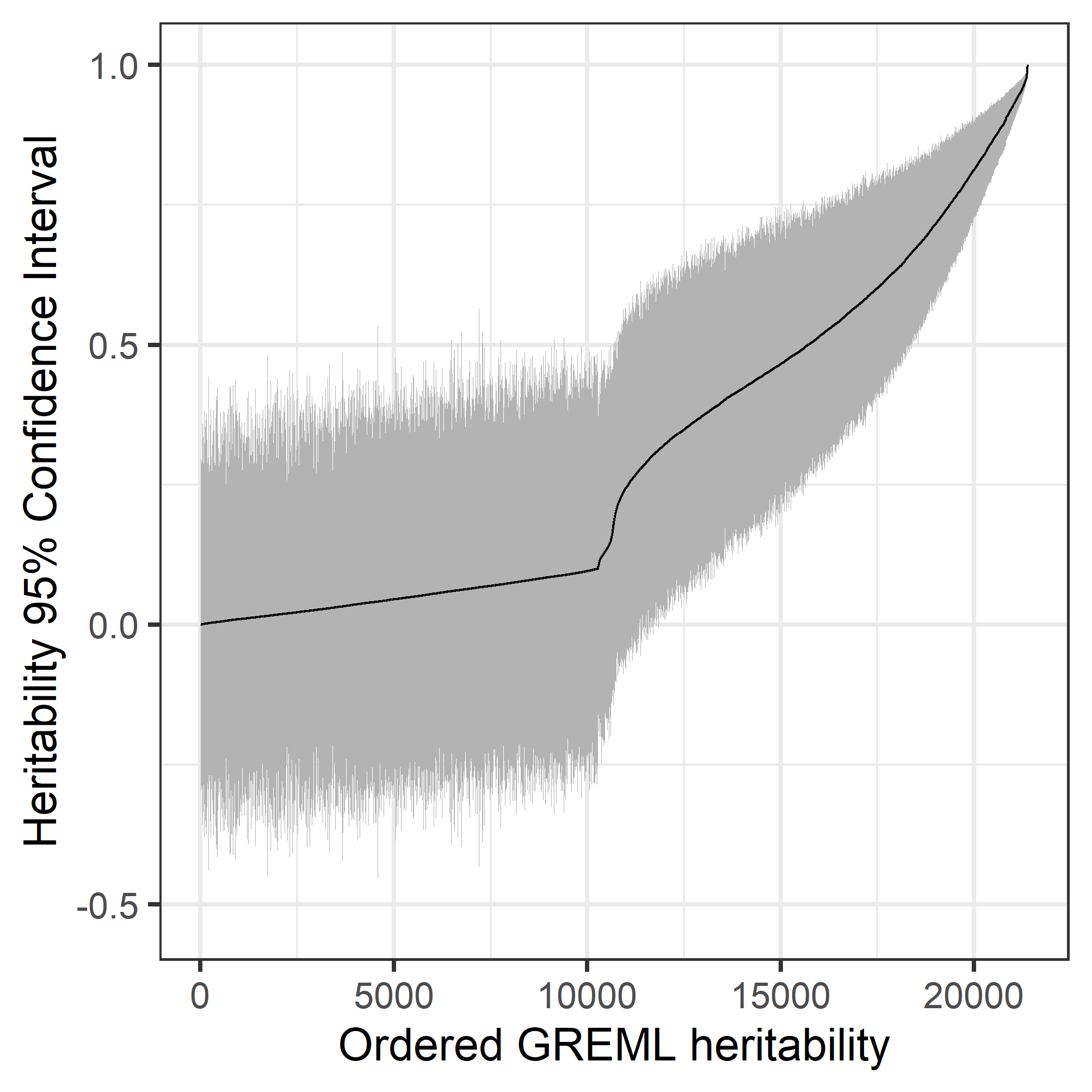

**Supplementary Figure S1. Single-step GREML.** Estimates with confidence intervals: Heritability estimates using the single-step GREML method were ordered and plotted (black line). The 95% confidence estimates were also calculated (grey lines). Based on randomized permutation testing, a threshold was calculated using a fixed type-I error rate (α=0.01) (red dashed line).

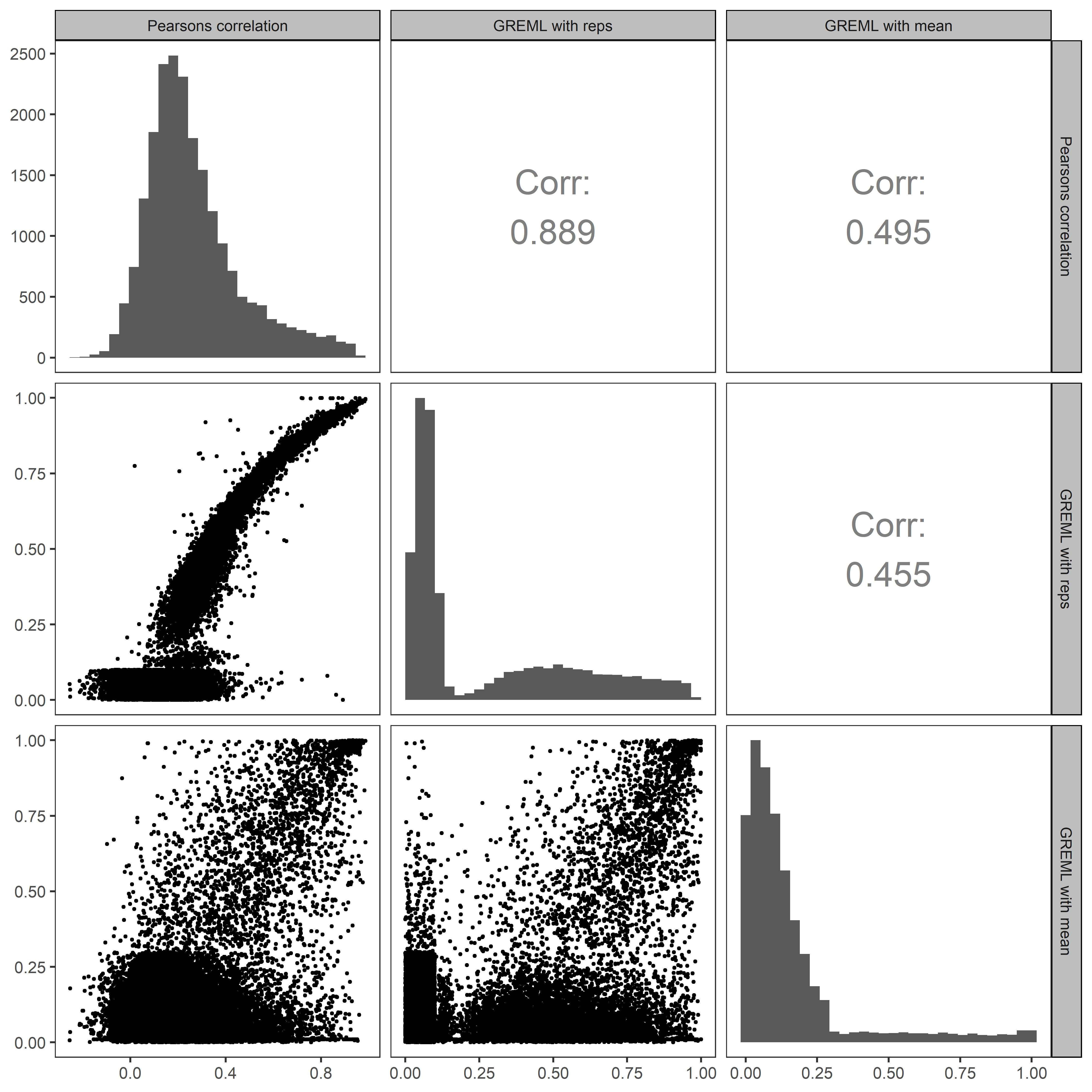

**Supplementary Figure S2. Comparison of Heritability Calculation Methods for Salt-Stress**. Pairwise correlation between repeatability (Pearson), single-step GREML (with replicates), and two-step GREML (using the genotypic mean) for the salt-stress condition.

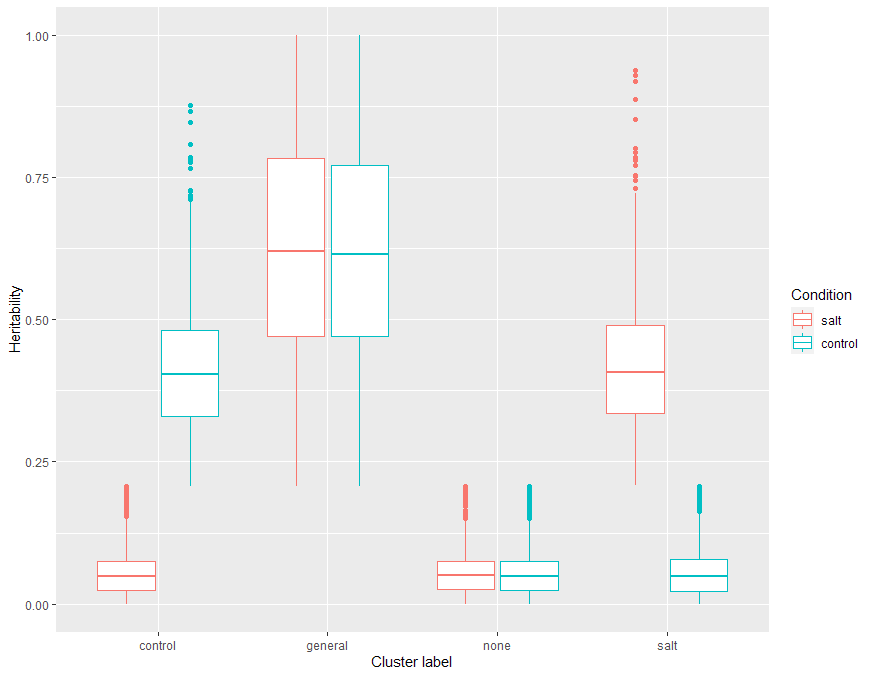

**Supplementary Figure S3. Condition-specific Heritabilities.** Genes were organized into groups of significant heritability in salt treatment, control, both or neither (none). The distribution of GREML calculated heritability, *h*^2^, in each group is shown.

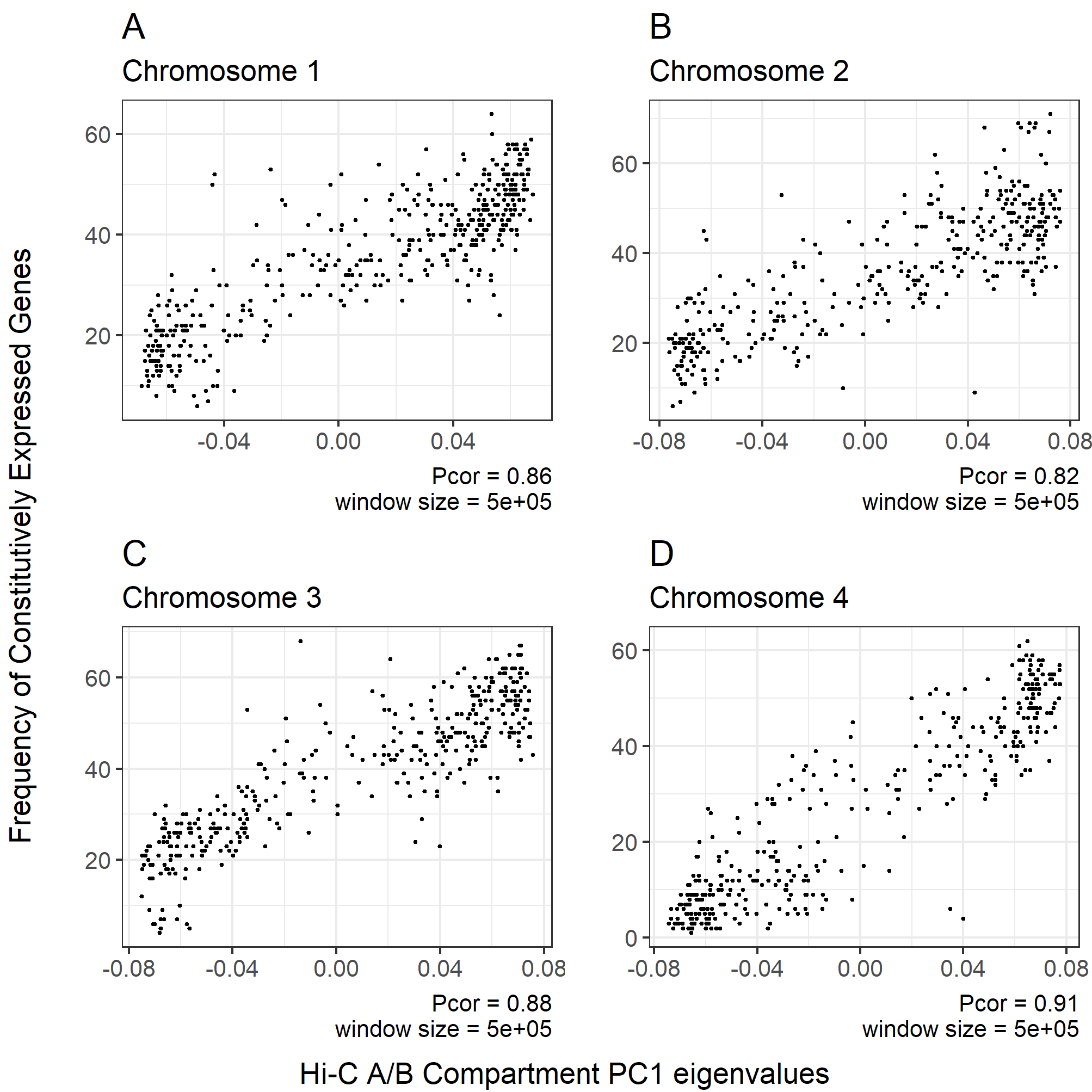

**Supplementary Figure S4. Comparison of Hi-C and Gene Expression: Chromosomes 1 (A), 2 (B), 3 (C), and 4 (D).** Eigenvectors of Hi-C A/B compartments are compared with the frequency of genes with constitutive expression (>95% of samples) for each chromosome using a fixed window size of 0.5 Mb.

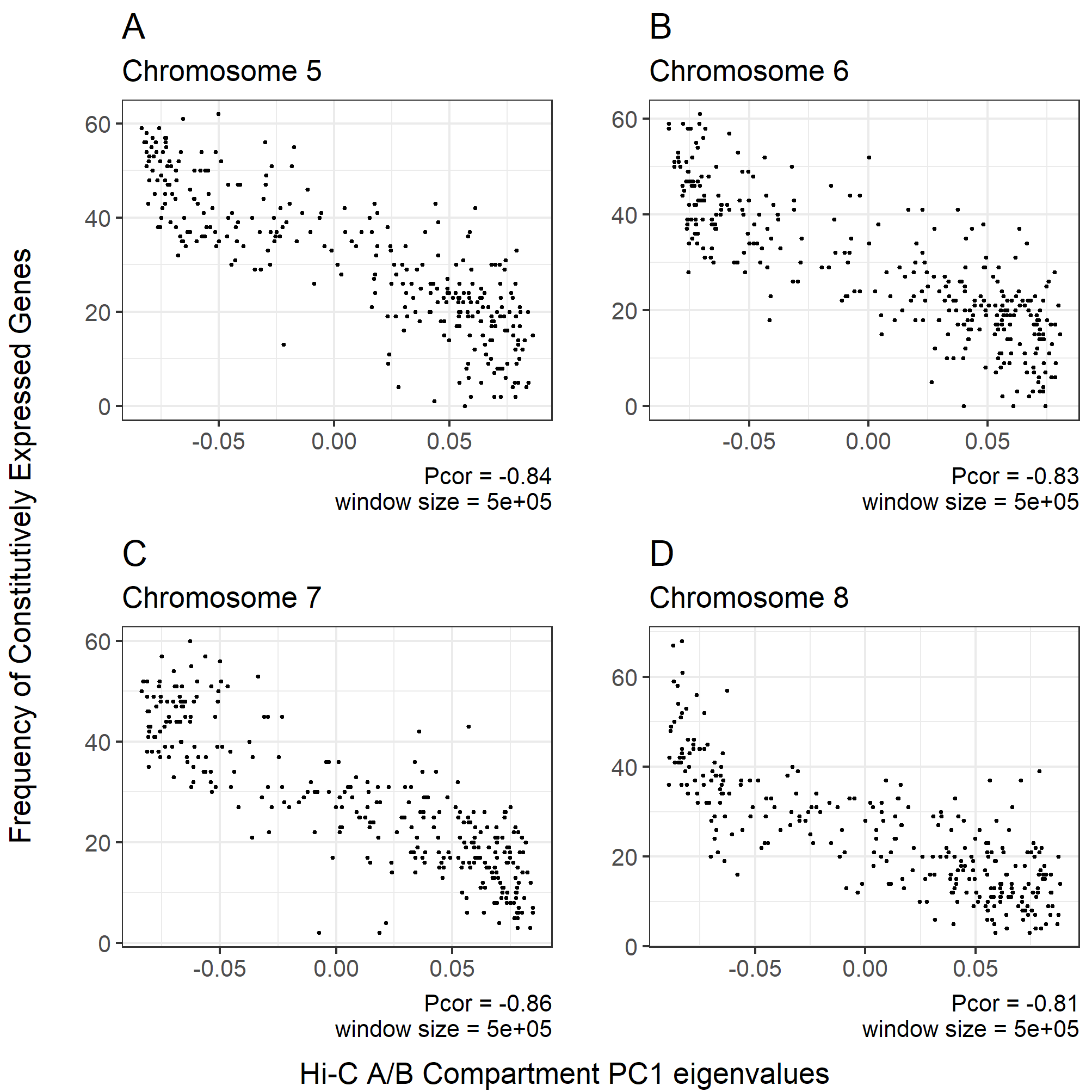

**Supplementary Figure S5. Comparison of Hi-C and Gene Expression: Chromosomes 5 (A), 6 (B), 7 (C), and 8 (D).** Eigenvectors of Hi-C A/B compartments are compared with the frequency of genes with constitutive expression (>95% of samples) for each chromosome using a fixed window size of 0.5 Mb.

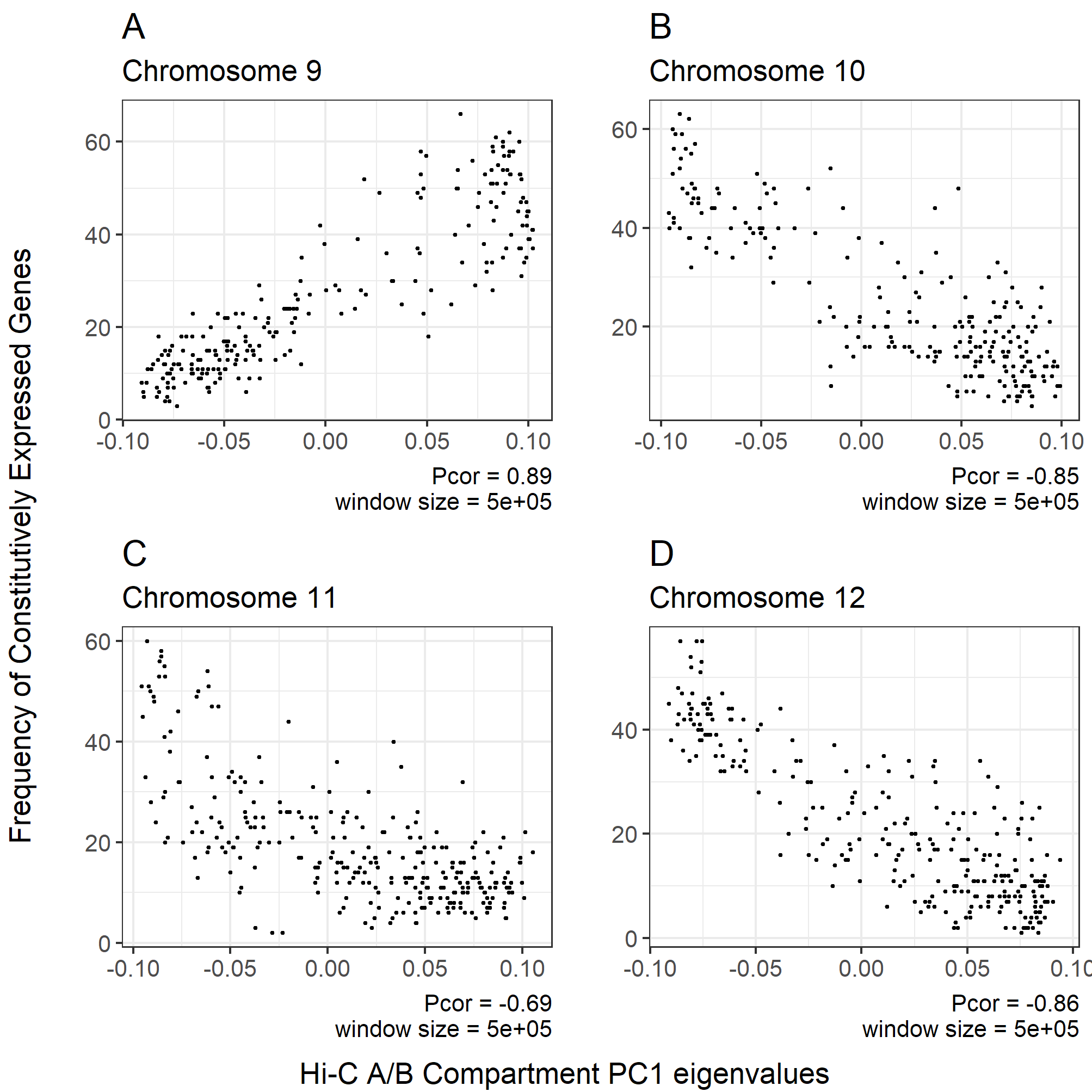

**Supplementary Figure S6. Comparison of Hi-C and Gene Expression: Chromosomes 9 (A), 10 (B), 11 (C), and 12 (D).** Eigenvectors of Hi-C A/B compartments are compared with the frequency of genes with constitutive expression (>95% of samples) for each chromosome using a fixed window size of 0.5 Mb.

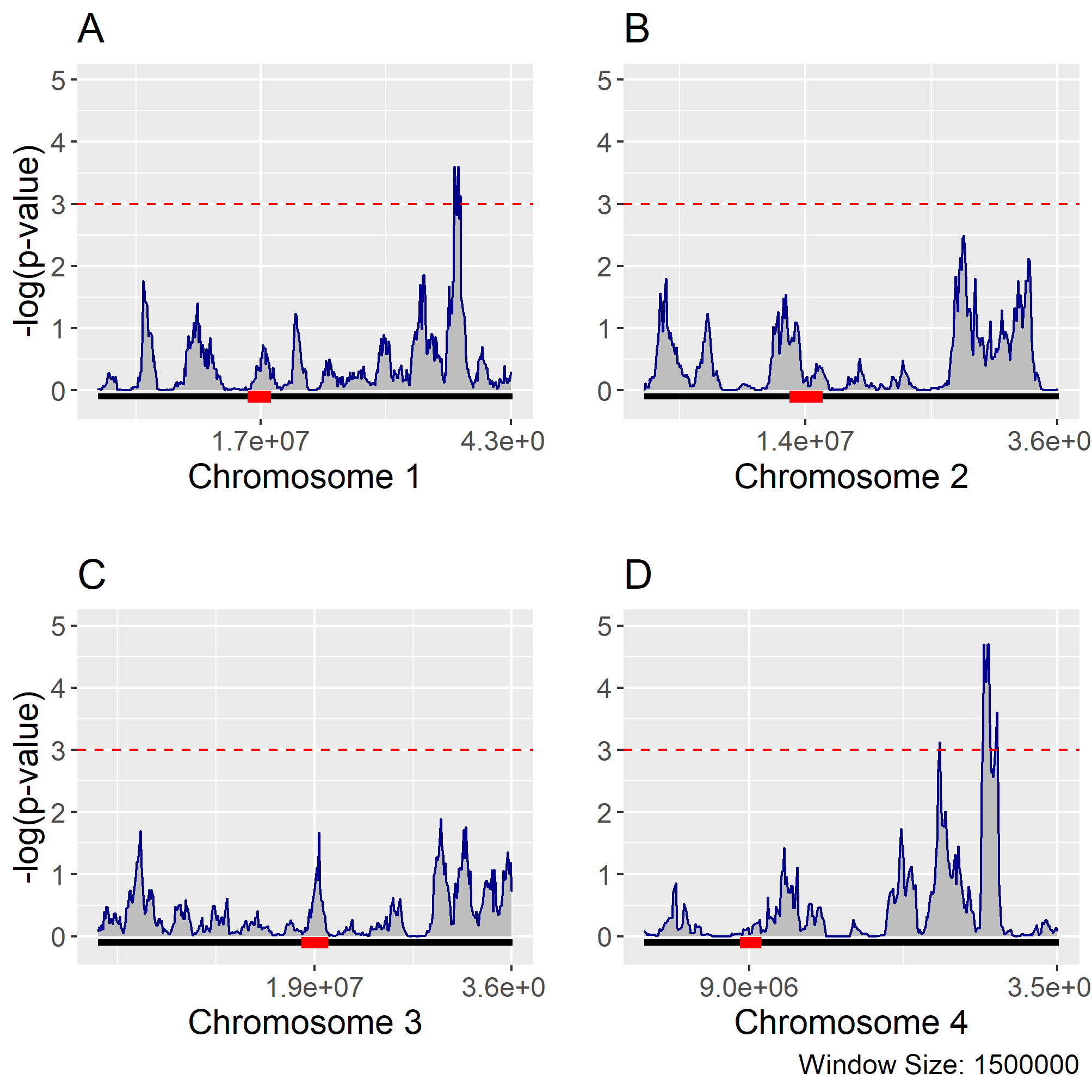

**Supplementary Figure S7. Salt-specific Heritability Enrichment: Chromosomes 1 (A), 2 (B), 3 (C), and 4 (D).** Using a sliding window size of 1.5 Mb at 100 Kb intervals, chromosomes were tested for enrichment of genes with salt-specific heritability using all genes with heritable expression (salt-specific, optimal-specific, and general) as the null distribution. P-values were adjusted for multiple-testing using a permutation based approach. Using a critical value of 0.001, significant windows enriched for salt-specific heritability were identified on chromosomes 1, 4, 6, and 8.

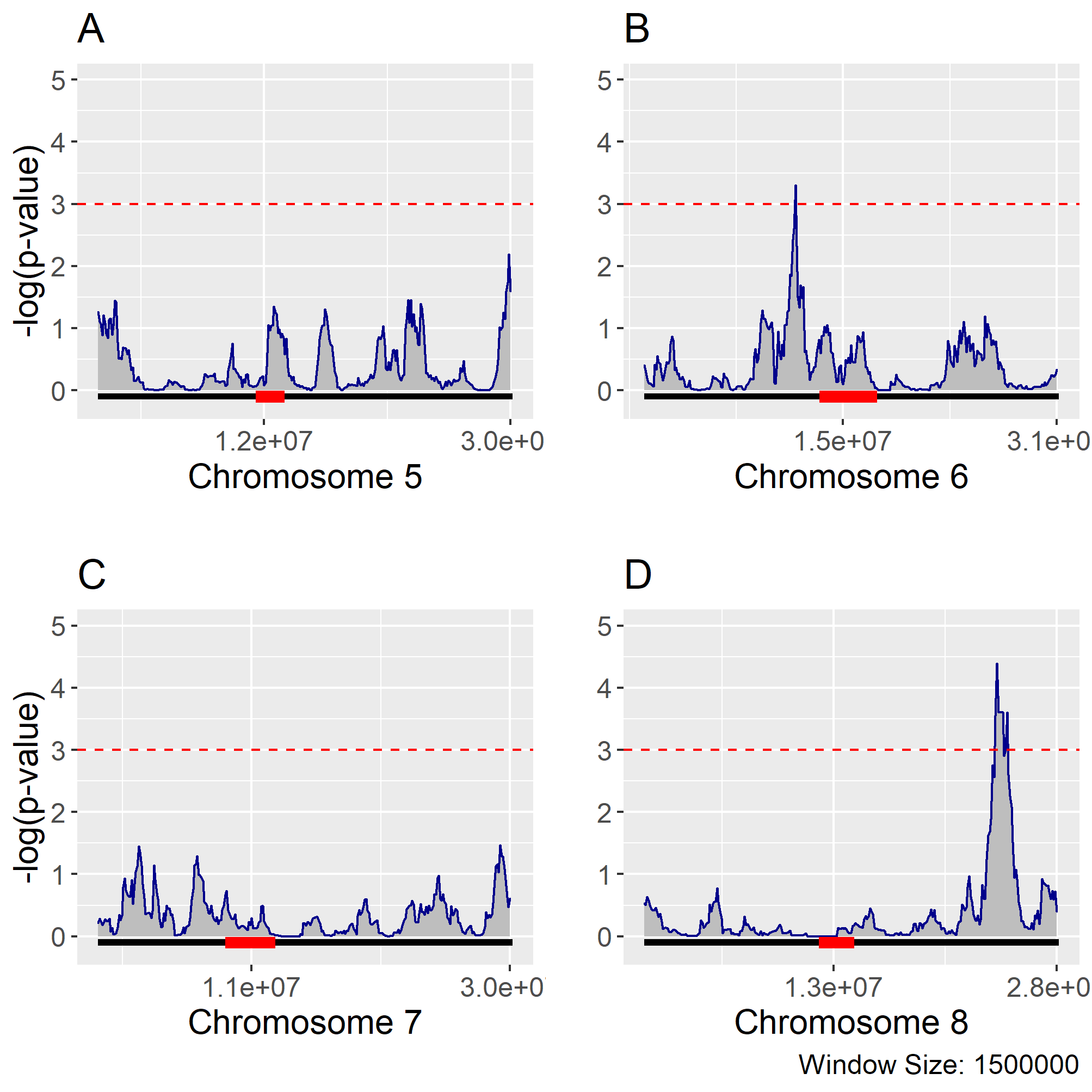

**Supplementary Figure S8. Salt-specific Heritability Enrichment: Chromosome 5 (A), 6 (B), 7 (C), and 8 (D).** Using a sliding window size of 1.5 Mb at 100 Kb intervals, chromosomes were tested for enrichment of genes with salt-specific heritability using all genes with heritable expression (salt-specific, optimal-specific, and general) as the null distribution. P-values were adjusted for multiple-testing using a permutation based approach. Using a critical value of 0.001, significant windows enriched for salt-specific heritability were identified on chromosomes 1, 4, 6, and 8.

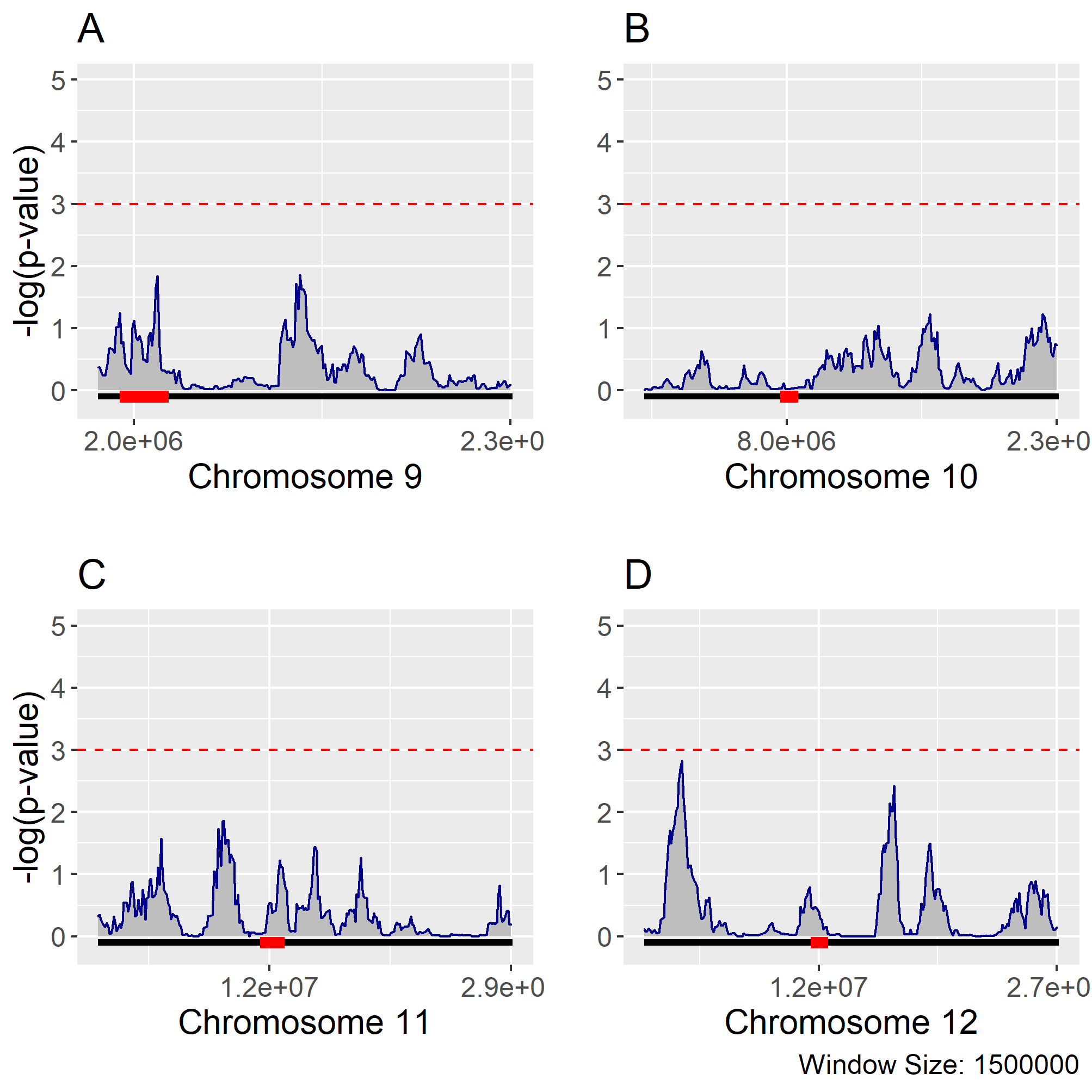

**Supplementary Figure S9. Salt-specific Heritability Enrichment: Chromosome 9 (A), 10 (B), 11 (C), and 12 (D).** Genome enrichment for salt-specific heritable expression: Using a sliding window size of 1.5 Mb at 100 Kb intervals, chromosomes were tested for enrichment of genes with salt-specific heritability using all genes with heritable expression (salt-specific, optimal-specific, and general) as the null distribution. P-values were adjusted for multiple-testing using a permutation based approach. Using a critical value of 0.001, significant windows enriched for salt-specific heritability were identified on chromosomes 1, 4, 6, and 8.

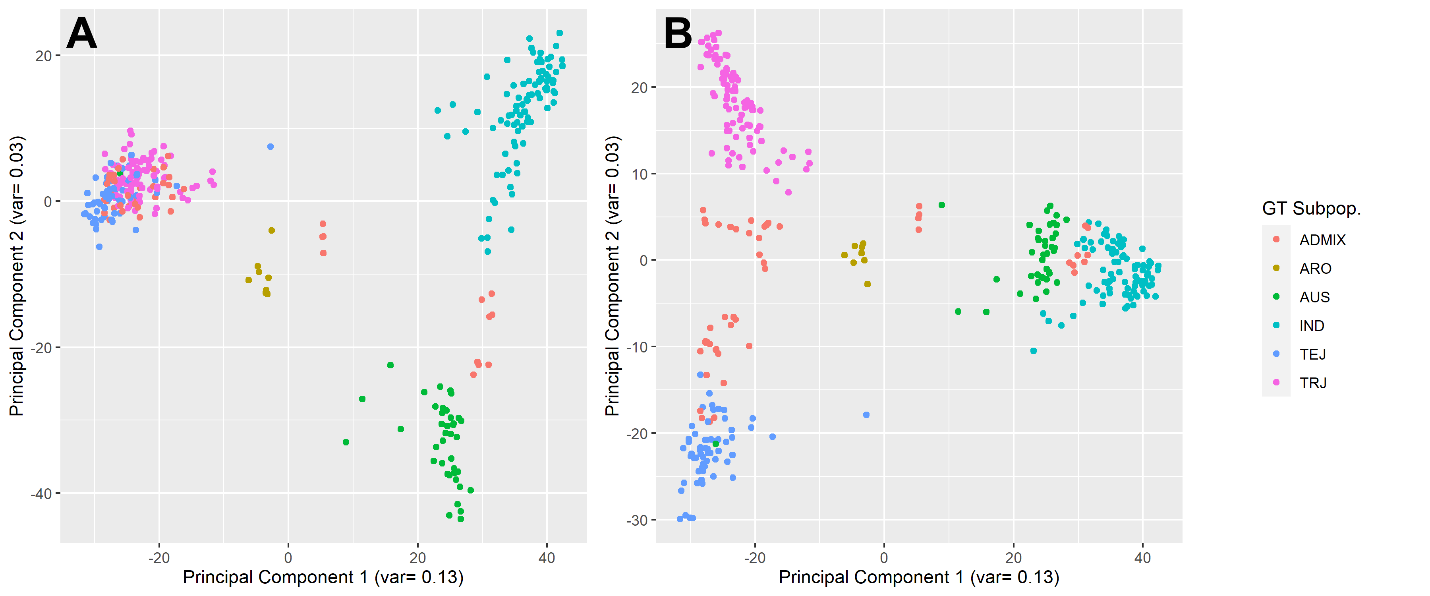

**Supplemental Figure S10. PCA Using Ordinal Categorical Gene Expression.** Scatterplots of the top three principal components calculated from a gene expression matrix including all RNAseq samples encoded as zero, one, or two based on whether TPMs were zero, low, or high respectively (based on the bimodal distribution). The top three components closely match principal components calculated using genotypes instead of gene expression (see Figure S11).

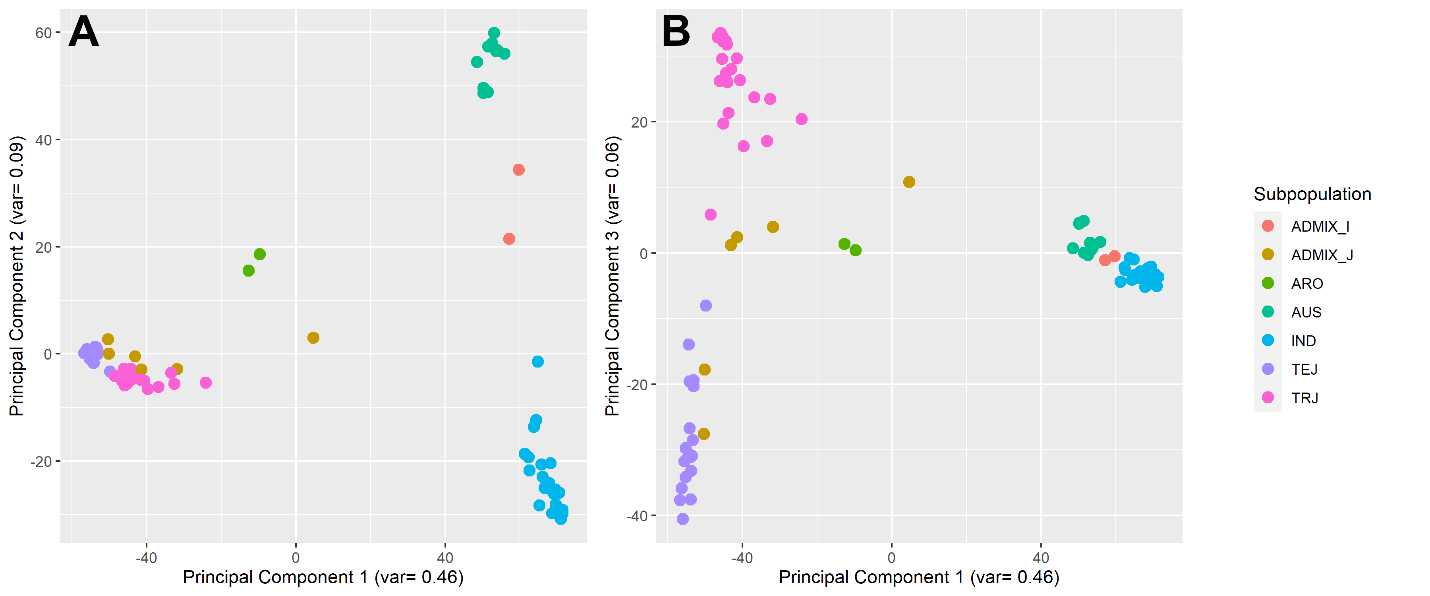

**Supplemental Figure S11. Rice Diversity Panel 1 Population Structure.** Scatterplots of the top three principal components calculated from genotypes of the 84 individuals subset from the RDP1 panel match domestication history in rice. Component one (A, B) corresponds to variance between japonica (TEJ and TRJ) and indica (IND) varieties. Component two (A) captures variance between indica and aus varieties. Component three (B) captures variance between temperate (TEJ) and tropical (TRJ) japonica varieties. Hybrid varieties (ADMIX_I and ADMIX_J) and aromatic (ARO) varieties occupied areas between major subspecies clusters

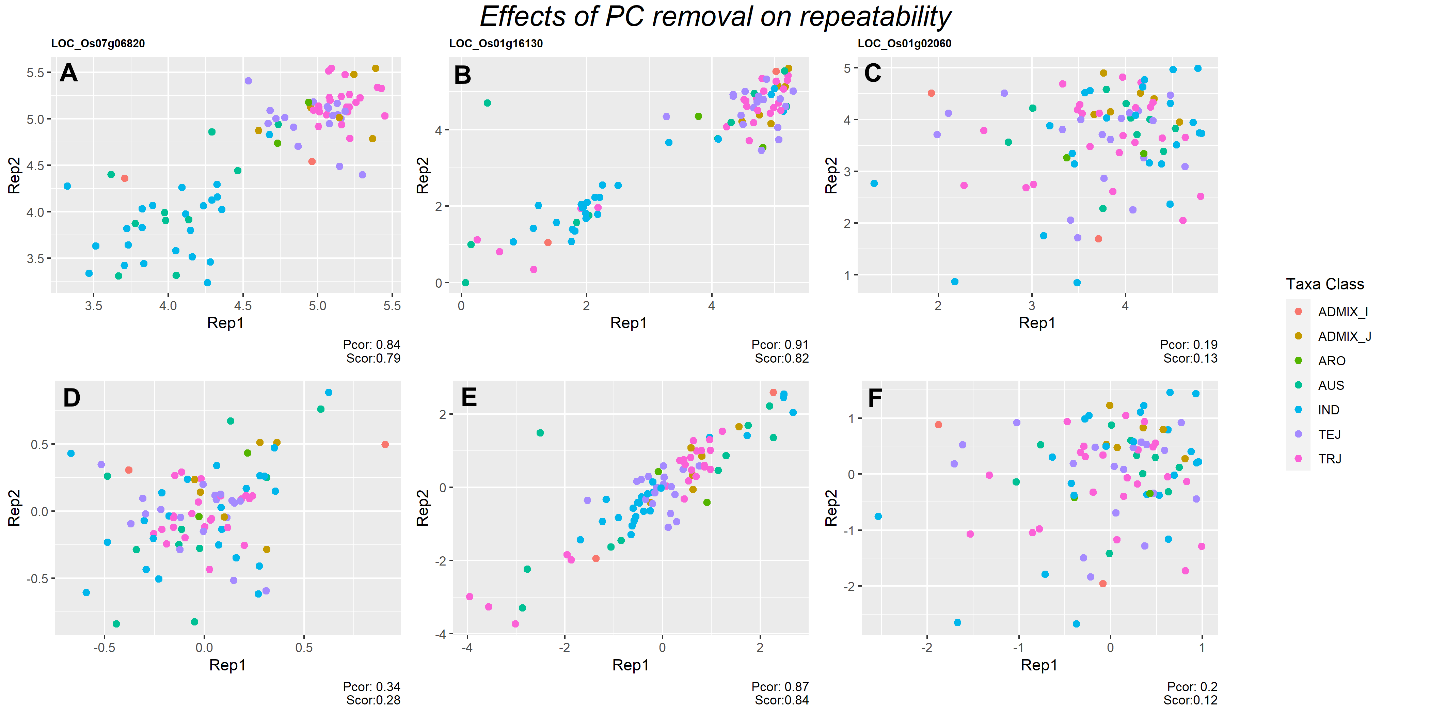

**Supplemental Figure S12. Adjustment for population structure**: Scatterplots of pair-wise unadjusted gene expression between replicate genotypes for three selected genes (A-C) demonstrate how population structure can cause gene expression to form distinct clusters and have high correlation (repeatability) between replicates. When this structure is removed, the repeatability is reduced for genes with expression primarily explained by those structures (D), but is largely unaffected for genes with non-structural variance in gene expression (E-F).
